# Inbreeding is associated with shorter early-life telomere length in a wild passerine

**DOI:** 10.1101/2021.10.10.463797

**Authors:** Michael Le Pepke, Alina K. Niskanen, Thomas Kvalnes, Winnie Boner, Bernt-Erik Sæther, Thor Harald Ringsby, Henrik Jensen

**Affiliations:** Centre for Biodiversity Dynamics (CBD), Department of Biology, Norwegian University of Science and Technology (NTNU), Trondheim, Norway; Ecology and Genetics Research Unit, University of Oulu, Oulu, Finland; Institute of Biodiversity, Animal Health and Comparative Medicine (IBAHCM), University of Glasgow, UK

**Author notes:** Correspondence: Michael Le Pepke,. Joint senior authors (Thor Harald Ringsby, Henrik Jensen).

**Keywords:** biomarker, conservation physiology, heterosis, inbreeding depression, SNP, telomere dynamics

## Abstract

Inbreeding can have negative effects on survival and reproduction, which may be of conservation concern in small and isolated populations. However, the physiological mechanisms underlying inbreeding depression are not well-known. The length of telomeres, the DNA sequences protecting chromosome ends, has been associated with health or fitness in several species. We investigated effects of inbreeding on early-life telomere length in two small island populations of wild house sparrows (*Passer domesticus*) known to be affected by inbreeding depression. Using genomic and pedigree-based measures of inbreeding we found that inbred nestling house sparrows have shorter telomeres. This negative effect of inbreeding on telomere length may have been complemented by a heterosis effect resulting in longer telomeres in individuals that were less inbred than the population average. Furthermore, we found some evidence of stronger effects of inbreeding on telomere length in males than females. Thus, telomere length may reveal subtle costs of inbreeding in the wild and demonstrate a route by which inbreeding negatively impacts the physiological state of an organism already at early life-history stages.

## INTRODUCTION

Inbreeding may have significant detrimental effects on survival, reproduction, and resistance to disease and other stressors in wild populations (Keller & Waller, 2002). Such decline in fitness resulting from an increase in genome-wide homozygosity is known as inbreeding depression (Charlesworth & Willis, 2009) and is of major concern in small and isolated populations, in particular of endangered species (Bozzuto, Biebach, Muff, Ives, & Keller, 2019; Harrisson et al., 2019; Hedrick & Kalinowski, 2000). Increased homozygosity can lead to reduced fitness due to expression of deleterious recessive alleles (“dominance hypothesis”) or increased homozygosity at loci with heterozygote advantage (“overdominance hypothesis”, Charlesworth & Willis, 2009). Regardless of the genetic basis for inbreeding depression, it is difficult to identify and quantify the physiological mechanisms underlying the fitness costs of inbreeding (Fox & Reed, 2011; Kristensen, Pedersen, Vermeulen, & Loeschcke, 2010; Losdat, Arcese, Sampson, Villar, & Reid, 2016).

Telomeres are short DNA tandem repeats that are found at the tips of most eukaryotic chromosomes (Blackburn & Gall, 1978; Červenák, Sepšiová, Nosek, & Tomáška, 2021). Telomeres shorten during cell division (Harley, Futcher, & Greider, 1990), but may also shorten due to several other reasons including physiological processes generating oxidative stress (Barnes, Fouquerel, & Opresko, 2019; Monaghan & Ozanne, 2018; Reichert & Stier, 2017; von Zglinicki, 2002). The high guanine content of telomeres (50%) makes them particularly vulnerable to oxidative stress (Kawanishi & Oikawa, 2004). Short telomeres can trigger apoptosis and telomere attrition is considered a hallmark of aging (López-Otín, Blasco, Partridge, Serrano, & Kroemer, 2013), although the causal involvement of telomere shortening in organismal senescence is not well understood (Simons, 2015). However, telomere length (TL) may reflect the cumulative stress experienced by an individual (Bateson, 2016; Monaghan, 2014), and TL or TL shortening are associated with health or fitness in several species (Barrett, Burke, Hammers, Komdeur, & Richardson, 2013; Chatelain, Drobniak, & Szulkin, 2020; Froy et al., 2021; Heidinger, Kucera, Kittilson, & Westneat, 2021; Wilbourn et al., 2018). Thus, TL is increasingly used as a biomarker of somatic integrity in studies of physiological or evolutionary ecology (Bateson & Poirier, 2019; Haussmann, 2010; Pepper, Bateson, & Nettle, 2018; Young, 2018).

Inbreeding depression can be caused by reduced immune response (Charpentier, Williams, & Drea, 2008; Reid, Arcese, & Keller, 2003) and higher maintenance metabolism (Ketola & Kotiaho, 2009), which increases oxidative stress (de Boer et al., 2018a; Okada, Blount, Sharma, Snook, & Hosken, 2011). Thus, inbred individuals may experience higher levels of oxidative stress (Kristensen, Sørensen, Kruhøffer, Pedersen, & Loeschcke, 2005; Pedersen et al., 2008) and thus have shorter telomeres (von Zglinicki, 2002). We therefore hypothesize that TL could provide an integrative measure of the somatic costs associated with inbreeding depression in wild populations, with inbred individuals having shorter telomeres than outbred individuals. However, the few studies investigating associations between inbreeding and TL have found equivocal results. In line with our expectations, Bebbington et al. (2016) found that homozygosity was negatively associated with TL in wild Seychelles warblers (*Acrocephalus sechellensis*) and Seluanov et al. (2008) reported that telomeres were shorter in inbred laboratory strains of Norway rats (*Rattus norvegicus*) in captivity compared to a single wild-caught rat. Many domesticated species are generally assumed to be more inbred than their wild counterparts (Bosse, Megens, Derks, de Cara, & Groenen, 2018; Moyers, Morrell, & McKay, 2018; Wiener & Wilkinson, 2011). However, several studies have found that telomeres were longer in inbred domesticated strains of laboratory mice (*Mus* spp. and *Peromyscus* spp., Hemann & Greider, 2000; Manning, Crossland, Dewey, & Van Zant, 2002; Seluanov et al., 2008), in domesticated strains of pearl millet (*Pennisetum glaucum*, Sridevi, Uma, Sivaramakrishnan, & Isola, 2002), in domesticated inbred chicken (*Gallus gallus*, O’Hare & Delany, 2009), and across several species of domesticated mammals (Pepke & Eisenberg, 2021) compared to non-domesticated species. However, there were no clear differences in TL between inbred and wild leporid strains (Forsyth, Elder, Shay, & Wright, 2005). Other studies found no association between pedigree-based inbreeding coefficients and TL or telomere attrition in humans (*Homo sapiens*, Mansour et al., 2011), wild sand lizards (*Lacerta agilis*, Olsson, Wapstra, & Friesen, 2018), or wild natterjack toads (*Epidalea calamita*, Sánchez-Montes et al., 2020). Becker et al. (2015) reported a weak non-significant but positive association between inbreeding and TL in wild white-throated dippers (*Cinclus cinclus*).

These contrasting results suggest that the telomere dynamics of captive, domesticated species living in a controlled environment may not be representative of wild, free-living populations (Chatelain et al., 2020; Pepke & Eisenberg, 2021; Weinstein & Ciszek, 2002). For instance, captive populations may be less vulnerable to inbreeding because inbreeding depression is greater under stressful environmental conditions (Fox & Reed, 2011; Reed, Briscoe, & Frankham, 2002). Furthermore, captivity may in itself provide conditions that change the telomere dynamics of the populations (Eisenberg, 2011), e.g. Hemann and Greider (2000) attributed the longer telomeres of inbred mice to effects of captive breeding and not inbreeding *per se*. For instance, TL shortening rates may increase during metabolically costly processes such as reproduction (Sudyka, Arct, Drobniak, Gustafsson, & Cichoń, 2019; Wood et al., 2021) and inbreeding may reduce fecundity (Keller & Waller, 2002). Such effects have been suggested to explain the observation of longer adult TL in some inbred domesticated species (Eisenberg, 2011), which could be resolved by measuring TL in early-life. Furthermore, most of the studies of domesticated animals compared TLs of different populations or species and their results may not be extrapolated to natural variation in TL and inbreeding levels within wild populations. Indeed, TL can vary considerably within species (Tricola et al., 2018) and across closely related species (Pepke, Ringsby, & Eisenberg, 2021) in the wild. Finally, it is not known if outbreeding could be accompanied by a heterosis effect (hybrid vigor, e.g. Charlesworth & Willis, 2009) acting on TL. For instance, the observed fitness benefits of outcrossing inbred populations (Frankham, 2015) could be reflected in TL restoration (Nuzhdin & Reiwitch, 2002; Ozawa et al., 2019).

In this study, we utilized a long-term metapopulation study to examine how inbreeding affects early-life TL in wild house sparrows (*Passer domesticus*). Inbreeding has been shown to reduce fitness components such as recruitment probability, adult lifespan, and both annual and lifetime reproductive success in this metapopulation (Billing et al., 2012; Jensen, Bremset, Ringsby, & Sæther, 2007; Niskanen et al., 2020), but the physiological effects underlying these phenomena remain unknown. We expect that inbred individuals will have shorter telomeres if TL is a general biomarker of somatic integrity and health (e.g. Bebbington et al., 2016; Boonekamp, Simons, Hemerik, & Verhulst, 2013; Wilbourn et al., 2018). The effects of inbreeding on TL might be sex-specific (Benton et al., 2018; Billing et al., 2012; de Boer et al., 2018a; de Boer, Eens, & Müller, 2018b) or depend on environmental conditions (Armbruster & Reed, 2005; Szulkin & Sheldon, 2007). However, TL is negatively associated with body size or growth rate within many species (Monaghan & Ozanne, 2018; Ringsby et al., 2015) and may change with age (Hall et al., 2004; Remot et al., 2021) or vary between sexes (Barrett & Richardson, 2011; Remot et al., 2020) and habitat quality (Angelier, Vleck, Holberton, & Marra, 2013; McLennan et al., 2021; Wilbourn et al., 2017). We therefore account for body size (measured as tarsus length), age, sex, and habitat type, and test for an interaction between inbreeding levels and sex or habitat type, when investigating the association between TL and inbreeding. We use three different measures of inbreeding; marker-based estimates (*n*=371) which are better at capturing homozygosity and inbreeding caused by distant ancestors not included in a pedigree, and pedigree-based estimates (Kardos, Taylor, Ellegren, Luikart, & Allendorf, 2016) for which larger samples size may be obtained from long-term field studies (*n*=1195). Finally, to investigate a potential heterosis effect on TL, we test if the association between TL and inbreeding is different among outbred and inbred individuals.

## MATERIAL AND METHODS

### Study system

This study was conducted in two natural populations of house sparrows in northern Norway. On the island of Hestmannøy (66°33’N, 12°50‘E), the sparrows live around dairy farms, where they nest inside barns in cavities or nest boxes. The island is characterized by cultivated grassland, mountains, forest, and heathland. On the island of Træna (66°30’N, 12°05‘E), 34 km further from the mainland, the sparrows live in gardens of a small human settlement and nest in nest boxes. This island is dominated by heathland, sparse forest, and gardens. The natural breeding environment for house sparrows is human habitation (Hanson, Mathews, Hauber, & Martin, 2020) and they have evolved their commensal relationship with humans for millennia (Ravinet et al., 2018). While human presence or farming provide the natural basis of existence for house sparrows (Ringsby, Sæther, Jensen, & Engen, 2006), demographic characteristics, breeding densities, and inbreeding rates are comparable to other small isolated wild animal populations (Araya-Ajoy et al., 2021; Jensen et al., 2007; Niskanen et al., 2020). In the years 1994-2013 (on Hestmannøy) and 2004-2013 (on Træna), nestlings at the age of 5-14 days were ringed with a unique combination of color rings for identification. Nestlings were also blood sampled by brachial venipuncture, and tarsometatarsus (tarsus) was measured with slide calipers to the nearest 0.01 mm. Tarsus length is here used as an index of body size (Rising & Somers, 1989; Senar & Pascual, 1997). Blood samples (25 μL) were stored in 96% ethanol at room temperature in the field and at - 20°C in the laboratory until DNA extraction (described in Pepke et al., *submitted* 2021b). Birds that were resighted or recaptured in the year following hatching (i.e. from 1995-2014 on Hestmannøy and from 2005-2014 on Træna) were categorized as first-year survivors.

### Telomere length measurements

Relative erythrocyte telomere length (TL) was measured in DNA derived from whole blood samples (*n*=2746 nestlings) using the qPCR method (Cawthon, 2002) as described in Pepke et al. (*submitted* 2021a). For this study, we included only individuals with two known parents and at least two known grandparents, or for which genomic inbreeding coefficients could be estimated (described below), resulting in a sample size of *n*=1370 individuals (*n*=1161 from Hestmannøy and *n*=209 from Træna). TL was determined relative to the amount of a non-variable gene (GAPDH) and a reference sample (Criscuolo et al., 2009). All samples were randomized and run in triplicates on 96-well plates. All samples were processed within a few months by the same researcher (MLP) to reduce technical effects. Relative TL was computed using qBASE (Hellemans, Mortier, De Paepe, Speleman, & Vandesompele, 2007) while controlling for inter-run variation. All individual plate efficiencies were within 100±10% (see Pepke et al., *submitted* 2021a). Sex was determined by amplification of the CHD-gene as described in Jensen et al. (2007).

### Microsatellite pedigree construction

Microsatellite (MS) pedigrees (*n*=1857 individuals from Hestmannøy and *n*=342 from Træna including non-phenotyped ancestors) were constructed based on 13 polymorphic microsatellite markers using CERVUS 3.0 (Kalinowski, Taper, & Marshall, 2007) as described in Billing et al. (2012). Maximum pedigree depth was 13 generations. We calculated inbreeding coefficients (*F*_*PED*_), which estimate the expected proportion of an individual’s genome that is identical by descent (IBD), based on the MS pedigree for individuals with two known parents and at least two known grandparents (*n*=1057 from Hestmannøy and *n*=138 from Træna, Table 1) using the R package *pedigree* (Coster, 2012). We also selected a subset of individuals with at least two full ancestral generations (i.e. four known grandparents) to only include the most robust estimates of *F*_*PED*_ (*n*=313 from Hestmannøy and *n*=7 from Træna).

**Table 1:** Number of nestling house sparrows of each sex and in total with early-life telomere length and inbreeding coefficient measurements within each population (Hestmannøy and Træna) for each measure of inbreeding (microsatellite pedigree-based inbreeding coefficient [*F*_*PED*_], genomic inbreeding coefficient [*F*_*GRM*_], and runs-of-homozygosity [*F*_*ROH*_]). Number of individuals with at least two known full ancestral generations (gen.) are shown. Number of individuals with *F*_*GRM*_ values above and below the mean *F*_*GRM*_, which is used as a break point to differentiate individuals that were more and less inbred than average, respectively, are also shown.

| Island population: | Hestmannøy |  |  | Træna |  |  | Sum: |
| --- | --- | --- | --- | --- | --- | --- | --- |
|  | Males | Females | Sum: | Males | Females | Sum: |  |
| $F_{PED} (\geq 1.5 \text{ gen.})$ | 511 | 546 | 1057 | 78 | 60 | 138 | 1195 |
| $F_{PED} (\geq 2 \text{ full gen.})$ | 148 | 165 | 313 | 4 | 3 | 7 | 320 |
| $F_{GRM}$ | 140 | 135 | 275 | 49 | 47 | 96 | 371 |
| $F_{GRM} > 0.016$ | 43 | 63 | 106 | 26 | 32 | 58 | 164 |
| $F_{GRM} < 0.016$ | 97 | 72 | 169 | 23 | 15 | 38 | 207 |
| $F_{ROH}$ | 140 | 135 | 275 | 49 | 47 | 96 | 371 |

### Genomic inbreeding estimation

Starting from year 1997 (Hestmannøy) or 2004 (Træna), birds that survived until recruitment (*n*=275 from Hestmannøy and *n*=96 from Træna) were genotyped for 200,000 Single Nucleotide Polymorphisms (SNPs) as described in Lundregan et al. (2018). Two genomic inbreeding coefficients were then estimated using 118,810 autosomal SNPs not in strong linkage disequilibrium, as described in Niskanen et al. (2020). The weighted average homozygosity over all loci from the genomic relationship matrix (*F*_*GRM*_) was estimated for the whole metapopulation simultaneously using the GCTA software (Yang, Lee, Goddard, & Visscher, 2011). *F*_*GRM*_ gives more weight to homozygotes of the minor allele than of the major allele, and it is an estimate of the correlation between homologous genes of the two gametes of an individual relative to the current population (Yang et al., 2011). *F*_*GRM*_ can be negative if the probability that the two homologous genes of an individual are IBD is smaller than that of two homologous genes being drawn at random from the reference population (Wang, 2014; Yang et al., 2011). Thus, the individuals with the smallest estimates of *F*_*GRM*_ are expected to be outbred (hybrids) because of e.g. mating involving immigrants (Wang, 2014). The proportion of the genome within runs-of-homozygosity (*F*_*ROH*_ ranging from 0 to 1, McQuillan et al., 2008) was estimated using the PLINK software (Purcell et al., 2007). ROH arise through mating of individuals that are IBD, and may therefore be used to estimate inbreeding (Curik, Ferenčaković, & Sölkner, 2014).

### Statistical analyses

To test whether TL was affected by inbreeding, we fitted linear mixed models (LMMs) using the package *lme4* (Bates, Mächler, Bolker, & Walker, 2015) in R v. 3.6.3 (R Core Team, 2020). TL (response variable) was log_10_-transformed to conform to the assumption of normally distributed residuals and the models were fitted with a (continuous) fixed effect of one of the inbreeding coefficients (*F*_*PED*_ [*n*=1195], *F*_*PED*_ with at least two full generations known [*n*=320], *F*_*GRM*_ [*n*=371], or *F*_*ROH*_ [*n*=371], see Table 1 for sample size details). Since genomic estimators of inbreeding (*F*_*GRM*_ and *F*_*ROH*_) were only available for recruits (first-year survivors), we tested whether the relationship between TL and *F*_*PED*_ varied between survivors (“1”, *n*=206) and non-survivors (“0”, *n*=989) by including an interaction effect between *F*_*PED*_ and first-year survival. Tarsus length increases with nestling age, so tarsus length was age-corrected by taking the residuals from a regression of tarsus length on age and age squared. This allowed us to include both tarsus length and age in the models describing variation in TL. Thus, age-standardized tarsus length, fledgling age at sampling (in number of days), hatch day (ordinal date mean centered across years), population identity (categorical: Hestmannøy or Træna), and sex (categorical: male or female) were included as fixed effects in all models. We tested whether the effect of inbreeding on TL varied between sexes and populations by including two-way interaction terms between the inbreeding coefficient and sex or population identity. Random intercepts were fitted for year and brood identity to account for the non-independence of nestlings from the same year and brood. This also controls for within-brood effects of inbreeding levels (Olsson et al., 2018). We then tested whether the inclusion of the inbreeding coefficient and interaction terms improved the baseline model (without the inbreeding coefficient) by comparing the resulting 5 candidate models using Akaike’s information criterion corrected for small sample sizes (*AICc*, Akaike, 1973; Hurvich & Tsai, 1989). Akaike weights (*w*) and evidence ratios (*ER*) were calculated to determine the relative fit of models to the data (Burnham & Anderson, 2002). To investigate heterosis effects on TL, we tested if the slopes of the regression between *F*_*GRM*_ and TL differed between individuals that were more inbred than on average (*F*_*GRM*_ > mean *F*_*GRM*_) and individuals that were less inbred than average (*F*_*GRM*_ < mean *F*_*GRM*_). We did this by testing if the inclusion of a regression break point at the mean *F*_*GRM*_ improved the models by comparing the resulting 9 candidate models using AICc. Models were validated visually using diagnostic plots of residuals, and model parameters are from models refitted with restricted maximum likelihood (REML). Estimates are reported with standard errors (SE) and 95% confidence intervals (CI). Regression lines were visualized using *ggplot2* (Wickham, 2016).

## RESULTS

Individual MS pedigree-based inbreeding coefficients (*F*_*PED*_) varied from 0.000-0.250 (mean 0.007, 16.9% non-zero values). The highest ranked model explaining variation in TL included a negative effect of *F*_*PED*_, but only slightly improved the fit of the baseline model (Δ_*2:1*_*AICc*=0.8 [subscripts denote which ranked models are compared], *w*_1_=0.36, *ER*_1_=*w*_*1*_*/w*_*2*_=1.49, Table S1 in the supporting information). Thus, there was a tendency for TL to be shorter in more inbred sparrows (β_*F_PED*_=-0.169±0.101, CI=[-0.366, 0.028], *n*=1195, Fig. 1a and Table 2). The model ranked third (Δ_*3:1*_*AICc*=1.3) indicated that TL was less associated with *F*_*PED*_ in males than in females (β_*F_PED*sex[female]*_=-0.167±0.196, CI=[-0.549, 0.216]), while the model ranked fourth (Δ_*4*_*AICc*=1.9) indicated that TL was less associated with *F*_*PED*_ in the Hestmannøy population than in the Træna population (β_*F_PED*island[Hestmannøy]*_=0.115±0.314, CI=[-0.498, 0.728]). However, due to high uncertainty in these parameter estimates, these effects are not deemed reliable.

**Table 2:** Estimates, standard errors (SE), lower and upper 95% confidence intervals (CI) from the highest ranked model of *F*_*PED*_ predicting variation in early-life TL (*n*=1195, see Table S2 and Fig. 1a). The model included random intercepts for brood identity (ID) and year. Estimates with CIs not overlapping 0 are shown in italics.

| **Response variable: $\log_{10}(\text{TL})$** | ***Estimate*** | **SE** | **Lower CI** | **Upper CI** |
| --- | --- | --- | --- | --- |
| intercept | -3.1E-4 | 0.037 | -0.072 | 0.071 |
| inbreeding coefficient ($F_{PED}$) | -0.169 | 0.101 | -0.366 | 0.028 |
| tarsus length | -0.003 | 0.002 | -0.008 | 0.001 |
| sex [female] | -0.006 | 0.006 | -0.017 | 0.005 |
| *island identity [Hestmannøy]* | *0.025* | *0.012* | *0.001* | *0.049* |
| age | -0.003 | 0.002 | -0.007 | 0.001 |
| hatch day | -1.4E-4 | 1.5E-4 | -4.4E-4 | 1.7E-4 |
| $\sigma^2_{\text{brood ID}} (n=500)$ | 0.002 | | 0.001 | 0.003 |
| $\sigma^2_{\text{year}} (n=20)$ | 0.003 | | 0.001 | 0.006 |
| Marginal $R^2$ / Conditional $R^2$: 0.014 / 0.393 | | | | |

**Fig. 1:**
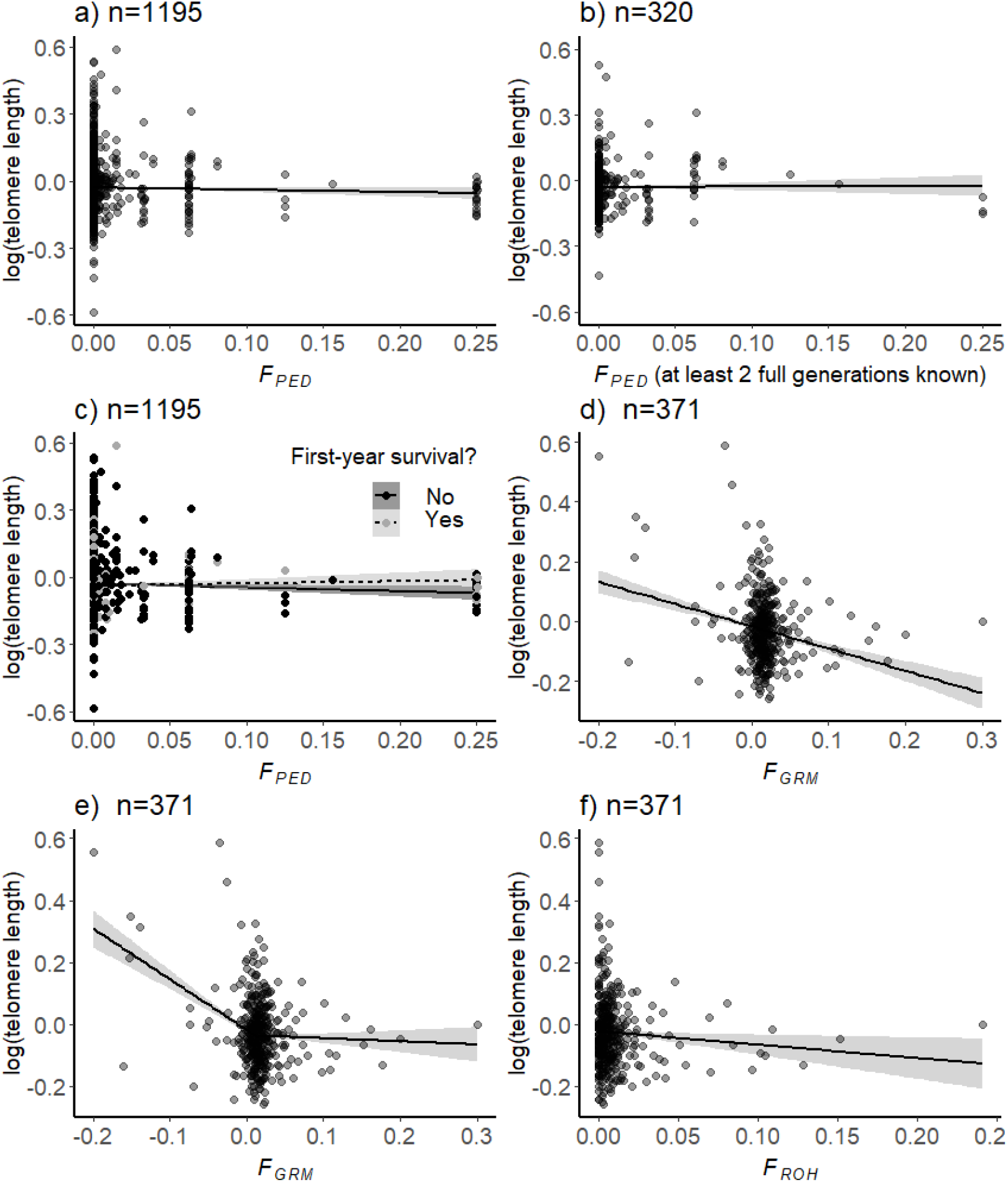
Associations between early-life telomere length (log_10_-transformed) and various individual measures of inbreeding in wild house sparrows: a) microsatellite pedigree-based inbreeding coefficient (*F*_*PED*_), b) *F*_*PED*_ for individuals with at least two full ancestral generations known, c) testing for an interaction effect between *F*_*PED*_ and first-year survival (survivors: *n*=206 in grey, dotted regression line; non-survivors: *n*=989 in black, solid regression line), d) genomic inbreeding coefficient *F*_*GRM*_, e) regression with a break point at the mean *F*_*GRM*_ (0.016), and f) runs-of-homozygosity *F*_*ROH*_. Black lines show the predicted effect of the inbreeding coefficient on TL from LMMs described in the text and the grey area shows 95% confidence intervals. Note that the y-axis is not scaled equally across panels and that color of points are graduated for visibility.

When only including individuals with at least 2 full ancestral generations known (33.8% non-zero values), the model with *F*_*PED*_ was ranked second (Δ_*2:1*_*AICc*=1.1, β_*F_PED*_=-0.205±0.198, CI=[-0.588, 0.189], *n*=320, Fig. 1b, Table S2-3) and the baseline model was highest ranked.

There was a tendency for the negative effect of *F*_*PED*_ on TL to be weaker in first-year survivors (*n*=206, mean TL=0.95±0.02, mean *F*_*PED*_=0.010±0.003) than in non-survivors (*n*=989, mean TL=0.97±0.01, mean *F*_*PED*_=0.007±0.001, β_*F_PED*first-year survival*_=0.304±0.201, CI=[-0.089, 0.697], *n*=1195, Fig. 1c, Table S4). This effect was uncertain with a CI overlapping zero. This suggests that the following analyses using genomic estimators of inbreeding in recruits were not biased towards stronger inbreeding effects in recruits.

Genomic inbreeding coefficient (*F*_*GRM*_) estimates varied from −0.200 to 0.300 (mean 0.016). The highest ranked model (Δ_*2:1*_*AICc*=2.1, Table S5) showed that TL was shorter in more inbred sparrows (β_*F_GRM*_=-1.517±0.293, CI=[-2.150, −0.920], *n*=371, Fig. 1d, and Table 3). In addition, the effect of *F*_*GRM*_ on TL was stronger in the Træna population (β_*F_GRM*island[Hestmannøy]*_=0.824±0.339, CI=[0.142, 1.529], Table 3) and in males (β_*F_GRM*sex[female]*_=0.644±0.314, CI=[0.034, 1.262], Table 3).

**Table 3:**
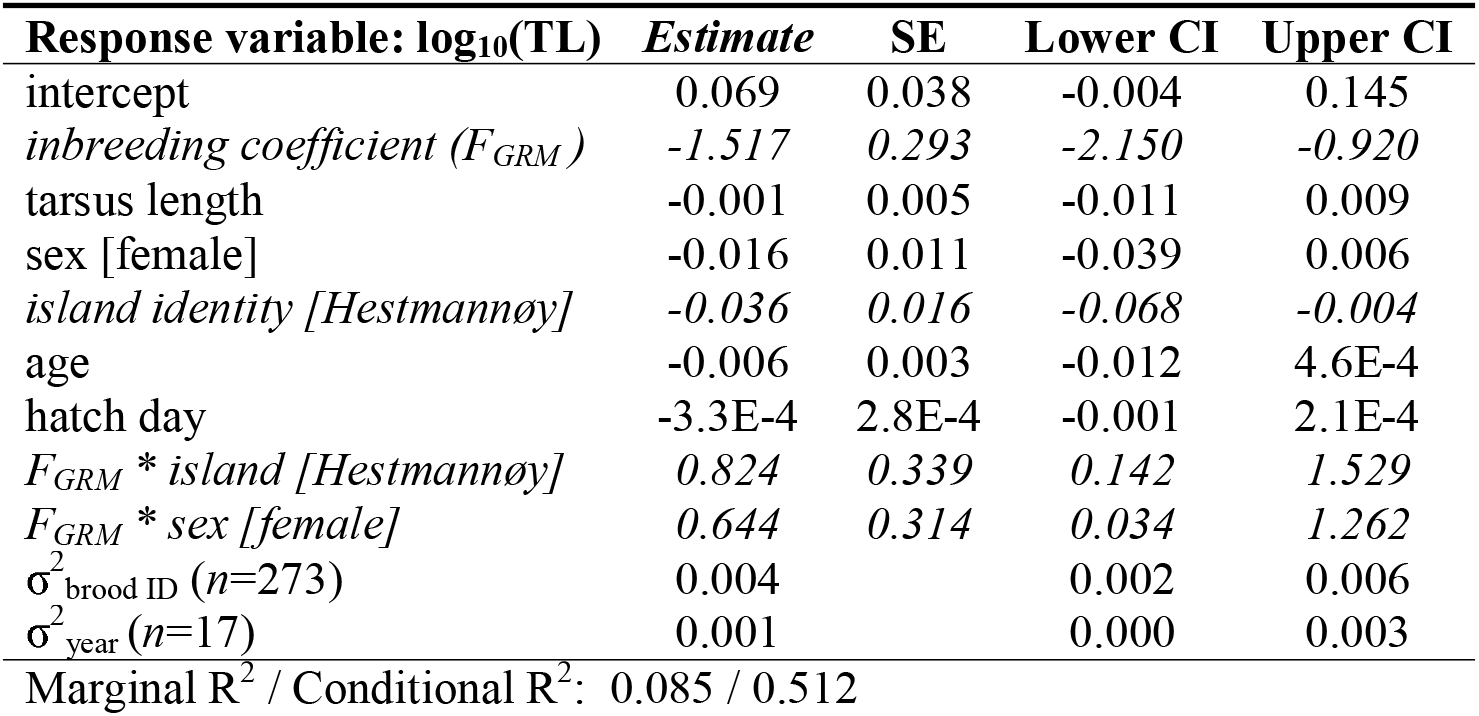
Estimates, standard errors (SE), lower and upper 95% confidence intervals (CI) from the highest ranked model of *F*_*GRM*_ predicting variation in early-life TL (*n*=371, see Table S6 and Fig. 1d).

| **Response variable: $\log_{10}(\text{TL})$** | ***Estimate*** | **SE** | **Lower CI** | **Upper CI** |
| --- | --- | --- | --- | --- |
| intercept | 0.069 | 0.038 | -0.004 | 0.145 |
| *inbreeding coefficient ($F_{GRM}$)* | *-1.517* | *0.293* | *-2.150* | *-0.920* |
| tarsus length | -0.001 | 0.005 | -0.011 | 0.009 |
| sex [female] | -0.016 | 0.011 | -0.039 | 0.006 |
| *island identity [Hestmannøy]* | *-0.036* | *0.016* | *-0.068* | *-0.004* |
| age | -0.006 | 0.003 | -0.012 | 4.6E-4 |
| hatch day | -3.3E-4 | 2.8E-4 | -0.001 | 2.1E-4 |
| $F_{GRM} * \text{island [Hestmannøy]}$ | *0.824* | *0.339* | *0.142* | *1.529* |
| $F_{GRM} * \text{sex [female]}$ | *0.644* | *0.314* | *0.034* | *1.262* |
| $\sigma^2_{\text{brood ID}} (n=273)$ | 0.004 | | 0.002 | 0.006 |
| $\sigma^2_{\text{year}} (n=17)$ | 0.001 | | 0.000 | 0.003 |
| Marginal $R^2$ / Conditional $R^2$: 0.085 / 0.512 | | | | |

Including a break point at the mean *F*_*GRM*_ improved the model compared to a model with no break point (comparing models without interaction terms which were ranked 8 and 5: Δ_*8:5*_*AICc*=4.5, see Table S6). The highest ranked model (Δ_*2:1*_*AICc*=3.1, Table S6) revealed a strong negative association between TL and *F*_*GRM*_ among individuals with *F*_*GRM*_<0.016 but no significant association among inbred individuals with *F*_*GRM*_>0.016 (Fig. 1e and Table 4). This indicates that a heterosis effect resulting in longer telomeres in outbred individuals may explain the negative association found between inbreeding and TL. This model also included an interaction term suggesting that this heterosis effect was stronger in the Træna population (Table 4).

**Table 4:** Estimates, standard errors (SE), lower and upper 95% confidence intervals (CI) from the highest ranked model from Table S7 including a break point at *F*_*GRM*_=0.016 (*n*=371, see also Table S1). These effects of *F*_*GRM*_ are shown in Fig. 1e.

| Response variable: $\log_{10}(TL)$ | Estimate | SE | Lower CI | Upper CI |
| --- | --- | --- | --- | --- |
| intercept | 0.021 | 0.037 | -0.051 | 0.095 |
| *inbreeding coefficient* ($F_{GRM} < 0.016$) | -2.177 | 0.372 | -3.051 | -1.379 |
| *inbreeding coefficient* ($F_{GRM} > 0.016$) | 0.189 | 0.498 | -0.780 | 1.153 |
| tarsus length | -0.001 | 0.005 | -0.011 | 0.008 |
| sex [female] | -0.006 | 0.010 | -0.027 | 0.014 |
| island identity [Hestmannøy] | -0.009 | 0.016 | -0.041 | 0.024 |
| age | -0.005 | 0.003 | -0.011 | 0.001 |
| hatch day | -3.7E-4 | 2.7E-4 | -0.001 | 1.5E-4 |
| $F_{GRM} < 0.016$ * island [Hestmannøy] | 1.562 | 0.465 | 0.610 | 2.576 |
| $F_{GRM} > 0.016$ * island [Hestmannøy] | -0.026 | 0.561 | -1.114 | 1.061 |
| $\sigma^2_{\text{brood ID}} (n=273)$ | 0.003 | | 0.001 | 0.005 |
| $\sigma^2_{\text{year}} (n=17)$ | 0.001 | | 0.000 | 0.003 |
| Marginal $R^2$ / Conditional $R^2$: 0.106 / 0.458 | | | | |

The runs-of-homozygosity inbreeding coefficient (*F*_*ROH*_) estimates varied from 0.000-0.240 (mean 0.010, 73% non-zero values). The best model provided evidence for a negative effect of *F*_*ROH*_ on TL (β_*F_ROH*_=-1.148±0.512, CI=[-2.144, −0.153], *n*=371, Fig. 1f, Table S7 and 5). This model also indicated that the negative effect of *F*_*ROH*_ tended to be stronger in males (β_*F_ROH*sex [female]*_=0.915±0.610, CI=[-0.270, 2.102]).

Overall, *F*_*PED*_ was not a good predictor of genomic estimators of inbreeding (Fig. S1a,c; Pearson’s *r*_*P*_=0.05, *n*=371), but its relationships with *F*_*GRM*_ and *F*_*ROH*_ were improved when including only individuals with at least two generations known (Fig. S1b,d; *r*_*P*_>0.30, *n*=59). *F*_*GRM*_ and *F*_*ROH*_ were strongly correlated (Fig. S1e,f; *r*_*P*_=0.7, *n*=371).

## DISCUSSION

We found evidence across multiple complementary measures of inbreeding that more inbred house sparrow nestlings had shorter telomeres (Fig. 1). Individual differences in TL are established early in life (Entringer, de Punder, Buss, & Wadhwa, 2018), are heritable (Dugdale & Richardson, 2018; Pepke et al., *submitted* 2021a), and are positively associated with fitness in some species (Heidinger et al., 2012; Wilbourn et al., 2018). Thus, short telomeres in more inbred individuals may therefore underpin a physiological basis of inbreeding depression in fitness components that has been found in this species (Billing et al., 2012; Jensen et al., 2007; Niskanen et al., 2020) and in other wild animal populations (Keller & Waller, 2002).

The effect of inbreeding on TL in house sparrows was negative across all measures of inbreeding, but strongest when using genomic levels of inbreeding (Fig. 1d-f), probably because they are better at capturing homozygosity causing inbreeding depression compared to using a pedigree-based estimator (Fig. 1a-c, Alemu et al., 2021; Huisman, Kruuk, Ellis, Clutton-Brock, & Pemberton, 2016; Kardos et al., 2016). Mating between full siblings or between parent and offspring (*F*=0.25) resulted in a severe reduction in (relative) TL of 58% (*F*_*GRM*_), 48% (*F*_*ROH*_) or 11% (*F*_*PED*_) compared to breeding between unrelated individuals (Tables 2, 3, and 5). TL may be under strong selection in natural populations (Voillemot et al., 2012). Consequently, strong inbreeding depression is expected for fitness components or traits that are under strong selection (Bérénos, Ellis, Pilkington, & Pemberton, 2016; DeRose & Roff, 1999), The analyses using genomic estimators of inbreeding were limited to recruited individuals, but the negative effect of inbreeding on TL may be even stronger if very inbred individuals, presumably with short telomeres, do not survive their first year and were thus excluded from our analyses (Jensen et al., 2007; Wilbourn et al., 2018). There was a tendency for such an effect when using pedigree-based levels of inbreeding (Fig. 1c and Table S4). We also found some evidence that inbreeding had stronger negative effects on TL in males than females (Tables 3 and 5). Such sex-specific effects of inbreeding are known from other species (de Boer et al., 2018a; de Boer et al., 2018b; Janicke, Vellnow, Sarda, & David, 2013), but have rarely been observed early in life. There was a weak tendency for longer TL in males than females (Tables 2-5), which has been observed in similar house sparrow populations (Pepke et al., *submitted* 2021b). Thus, males may be better buffered against the effects of inbreeding on TL. However, no sex-specific differences in inbreeding depression were observed in adult sparrows across this study metapopulation (Niskanen et al., 2020).

**Table 5:** Estimates, standard errors (SE), lower and upper 95% confidence intervals (CI) from the highest ranked model from of *F*_*ROH*_ predicting variation in early-life TL (*n*=371, see Table S8 and Fig. 1f).

| Response variable: $\log_{10}(TL)$ | Estimate | SE | Lower CI | Upper CI |
| --- | --- | --- | --- | --- |
| intercept | 0.051 | 0.040 | -0.027 | 0.130 |
| *inbreeding coefficient* ($F_{ROH}$) | -1.148 | 0.512 | -2.144 | -0.153 |
| tarsus length | -0.001 | 0.005 | -0.011 | 0.010 |
| sex [female] | -0.018 | 0.012 | -0.041 | 0.005 |
| island identity [Hestmannøy] | -0.020 | 0.016 | -0.052 | 0.012 |
| age | -0.005 | 0.003 | -0.012 | 0.001 |
| hatch day | -2.9E-4 | 3.0E-4 | -0.001 | 2.9E-4 |
| $F_{ROH}$ * sex [female] | 0.915 | 0.610 | -0.270 | 2.102 |
| $\sigma^2_{\text{brood ID}} (n=273)$ | 0.006 | | 0.004 | 0.008 |
| $\sigma^2_{\text{year}} (n=17)$ | 0.002 | | 4.6E-4 | 0.004 |
| Marginal $R^2$ / Conditional $R^2$: 0.029 / 0.579 | | | | |

Increased inbreeding may be accompanied by population decline in small populations (Bozzuto et al., 2019; Chen, Cosgrove, Bowman, Fitzpatrick, & Clark, 2016; Feng et al., 2019), which can drive populations to extinction (O’Grady et al., 2006; Saccheri et al., 1998; Wright, Tregenza, & Hosken, 2007). Niskanen et al. (2020) showed that inbreeding depression in adult sparrows in our study system varied little across years or across the different island environments inhabited by these house sparrows. Hence, the strength of inbreeding depression is similar between populations, but due to harboring more inbred individuals, the relative effect is stronger in smaller populations (Niskanen et al., 2020). Small declining populations may be characterized by gradual population-wide and trans-generational telomere erosion. For instance, Dupoué et al. (2017) observed shorter TL along an extinction risk gradient in populations of common lizards (*Zootoca vivipara*) that are disappearing from low altitudes at their southern range limit, presumably due to climate warming (Sinervo et al., 2010). Combined, these results suggest that TL may represent a potential physiological biomarker or molecular tool in conservation genetics addressing the viability of some small animal populations (Bebbington et al., 2016; Bergman et al., 2019; Dupoué et al., 2017; Madliger, Franklin, Love, & Cooke, 2020).

The negative effect of *F*_*GRM*_ on TL (Fig. 1d) was stronger among individuals that were less related than the average population (Fig. 1e). This suggests that longer telomeres in outbred individuals may partly be attributed to a general heterosis effect (Charlesworth & Willis, 2009) involving mating between immigrants and native individuals (Dickel et al., 2021; Ebert et al., 2002). In our study metapopulation, the proportion of dispersers among recruits can be high among the island populations (0.2 on average ranging from 0.0-1.0 across years and islands, Ranke et al., 2021; Saatoglu et al., 2021), and hence most islands are not strongly differentiated (Niskanen et al., 2020). We found that the negative effect of *F*_*GRM*_ on TL was stronger in the Træna population (Table 3-4). Træna is known to have a higher proportion of immigrants than Hestmannøy (Ranke et al., 2021), which may contribute to a stronger effect of heterosis in this population (Table 4). Furthermore, the gardens of Træna expose the sparrows to a different environment than the farms on Hestmannøy (Araya-Ajoy et al., 2019; Pärn, Ringsby, Jensen, & Sæther, 2012). Inbreeding depression is expected to have more severe consequences under environmental stress (Armbruster & Reed, 2005; Reed et al., 2002), such as harsh weather or competition (de Boer et al., 2018a; Fox & Reed, 2011; Marr, Arcese, Hochachka, Reid, & Keller, 2006). Telomeres shorten due to environmental stressors such as harsh abiotic conditions (Chatelain et al., 2020). We speculate that environmental differences between the habitats of the two sparrow populations may explain the exacerbated effects of inbreeding on TL in the Træna population. For instance, in juvenile Seychelles warblers a negative relationship between homozygosity and TL was found only in poor seasons, i.e. when food availability was low (Bebbington et al., 2016). In adult Seychelles warblers, the effect of homozygosity on TL was consistently negative across seasons, suggesting that the negative effects of inbreeding accumulate through life and are reflected in telomere erosion (Bebbington et al., 2016). Here, we showed that inbreeding manifests in TL already at the nestling stage in a similar wild passerine.

We measured TL in blood, thus it is possible that inbreeding or heterosis only affected telomeres in erythrocytes (Manning et al., 2002; Olsson, Geraghty, Wapstra, & Wilson, 2020). However, this is unlikely because TLs often correlate well across tissues within the organism (Daniali et al., 2013; Demanelis et al., 2020; Reichert, Criscuolo, Verinaud, Zahn, & Massemin, 2013), especially in early-life (Prowse & Greider, 1995). Although genomic inbreeding estimates were only available for first-year survivors, we may have avoided confounding effects of selective mortality of inbred individuals at much older ages by measuring TL already at the nestling stage (Hemmings, Slate, & Birkhead, 2012; Sánchez-Montes et al., 2020). Furthermore, since the mutation accumulation theory of senescence predicts that deleterious effects of inbreeding increase with age (Charlesworth & Hughes, 1996; Keller, Reid, & Arcese, 2008), we may expect that the effect on TL is persistent and potentially stronger in adult sparrows. Thus, future studies are required to investigate if inbreeding leads to persistently eroded TL throughout life, and if there are combined fitness consequences of any interaction between TL and inbreeding in wild populations. Even in the absence of a mechanism directly linking inbreeding and TL via the effects of oxidative stress (cf. the introduction), we may find inbred individuals to have short telomeres, because inbreeding impairs other physiological processes that affects both fitness and TL (Bebbington et al., 2016). Thus, the conflicting evidence in the literature of an effect of inbreeding on TL (reviewed in the introduction) suggests that an experimental procedure is needed to further elucidate the mechanisms underlying the correlation reported here (Manning et al., 2002), especially in wild populations.

In conclusion, the negative associations between inbreeding levels and TL found in this study suggest that TL may reveal subtle somatic costs of inbreeding in wild populations, and thereby demonstrates a potential route by which inbreeding negatively impacts the physiological state of an organism in early life. The observation of a potential heterosis effect on TL suggests that maintenance of dispersal within this metapopulation is important for mitigating the negative effects of inbreeding.

## ACKNOWLEDGEMENTS

We thank everyone contributing to the fieldwork, Randi Røsbak, Anna S. Båtnes, and Linn-Karina Selvik for performing DNA extractions for genotyping procedures, and Pat Monaghan for facilitating telomere analyses at the University of Glasgow and for commenting on an early version of this manuscript.

## DECLARATIONS

### Funding

This work was funded by the Research Council of Norway (274930 and 302619) and through its Centres of Excellence scheme (223257).

### Conflicts of interest

The authors have no conflicts of interest to declare.

### Availability of data and material

Data will be available on Dryad or another open data repository.

### Code availability

Not applicable.

### Authors’ contributions

MLP measured telomeres, analyzed data, and wrote the manuscript with contributions from all authors. WB supervised telomere measurements. HJ, AKN, and TK contributed to the genotype data processing, pedigree construction, and in designing statistical analyses. THR, BE-S, and HJ initiated the study system. THR, HJ, and TK contributed to the fieldwork.

### Ethics approval

Fieldwork was carried out in accordance with permits from the Ringing Centre at Stavanger Museum and the Norway Norwegian Animal Research Authority.

### Consent to participate

Not applicable.

### Consent for publication

Not applicable.

## Notes

### Competing Interest Statement

The authors have declared no competing interest.

