## Supporting information for "Inbreeding is associated with shorter early-life telomere length in a wild passerine"

### **AICc tables and model results of effects of *F_PED_* on telomere length**

**Table S1:** Linear mixed effects models (LMMs) of variation in early-life telomere length (*n*=1195) in two island populations of house sparrows. All models included random intercepts for brood identity and year. All models below are ranked by AICc, and number of degrees of freedom (df) and model weights (w) are shown.

| Model | | ∆AICc | | df | | w | |
| --- | --- | --- | --- | --- | --- | --- | --- |
| 1 | log(TL) = tarsus + sex + island + age + hatch day + *F_PED_* | | 0.0 | | 10 | | 0.36 |
| 2 | log(TL) = tarsus + sex + island + age + hatch day | | 0.8 | | 9 | | 0.24 |
| 3 | log(TL) = tarsus + sex + island + age + hatch day + *F_PED_* + *F_PED_******sex | | 1.3 | | 11 | | 0.19 |
| 4 | log(TL) = tarsus + sex + island + age + hatch day + *F_PED_* + *F_PED_**island | | 1.9 | | 11 | | 0.14 |
| 5 | log(TL) = tarsus + sex + island + age + hatch day + *F_PED_* + *F_PED_**sex + *F_PED_**island | | 3.2 | | 12 | | 0.07 |

**Table S2:** LMMs of variation in early-life telomere length (*n*=320 limited to individuals with at least 2 full ancestral generations known).

| Model | | ∆AICc | df | w |
| --- | --- | --- | --- | --- |
| 1 | log(TL) = tarsus + sex + island + age + hatch day | 0.0 | 9 | 0.47 |
| 2 | log(TL) = tarsus + sex + island + age + hatch day + *F_PED_* | 1.1 | 10 | 0.27 |
| 3 | log(TL) = tarsus + sex + island + age + hatch day + *F_PED_* + *F_PED_* ***** island | 2.8 | 11 | 0.12 |
| 4 | log(TL) = tarsus + sex + island + age + hatch day + *F_PED_* + *F_PED_* ***** sex | 3.1 | 11 | 0.10 |
| 5 | log(TL) = tarsus + sex + island + age + hatch day + *F_PED_* + *F_PED_* * sex + *F_PED_**island | 4.8 | 12 | 0.04 |

**Table S3:** Estimates, standard errors (SE), lower and upper 95% confidence intervals (CI) from the second highest ranked model in Table S2 (*∆_2_AICc*=1.1, *n*=320), which included the effect of *F_PED_*. The model included random intercepts for brood identity (ID) and year. The effect of *F_PED_* is shown in Fig. 1b.

| Response variable: log_10_(TL) | Estimate | SE | Lower CI | Upper CI |
| --- | --- | --- | --- | --- |
| intercept | 0.020 | 0.072 | -0.119 | 0.161 |
| inbreeding coefficient (*F_PED_*) | -0.205 | 0.198 | -0.588 | 0.189 |
| *tarsus length* | *-0.008* | *0.004* | *-0.016* | *-0.001* |
| *sex [female]* | *-0.024* | *0.011* | *-0.045* | *-0.004* |
| island identity [Hestmannøy] | 0.025 | 0.045 | -0.063 | 0.112 |
| *age* | *-0.010* | *0.003* | *-0.016* | *-0.003* |
| hatch day | -1.7E-4 | 3.0E-4 | -4.2E-4 | 0.001 |
| σ^2^_brood ID_ (*n*=147) | 0.002 |  | 0.001 | 0.004 |
| σ^2^_year_ (*n*=20) | 0.001 |  | 0.5E-4 | 0.003 |
| Marginal R^2^ / Conditional R^2^: 0.069 / 0.363 | | | | |

**Table S4:** Estimates, standard errors (SE), lower and upper 95% confidence intervals (CI) from a model including an interaction term between *F_PED_* and first-year survival (*n*=1195). The model included random intercepts for brood identity (ID) and year. The effect of *F_PED_* is shown in Fig. 1c.

| Response variable: log_10_(TL) | Estimate | SE | Lower CI | Upper CI |
| --- | --- | --- | --- | --- |
| intercept | -0.003 | 0.037 | -0.074 | 0.069 |
| *inbreeding coefficient (F_PED_)* | *-0.251* | *0.114* | *-0.474* | *-0.028* |
| tarsus length | -0.003 | 0.002 | -0.008 | 0.001 |
| sex [female] | -0.006 | 0.006 | -0.017 | 0.005 |
| *island identity [Hestmannøy]* | *0.026* | *0.012* | *0.002* | *0.049* |
| age | -0.003 | 0.002 | -0.007 | 0.001 |
| hatch day | -1.3E-4 | 1.6E-4 | -4.3E-4 | 1.7E-4 |
| first-year survival [1] | -0.002 | 0.008 | -0.017 | 0.014 |
| *F_PED_* * first-year survival [1] | 0.304 | 0.201 | -0.089 | 0.697 |
| σ^2^_brood ID_ (*n*=500) | 0.002 |  | 0.001 | 0.003 |
| σ^2^_year_ (*n*=20) | 0.003 |  | 0.001 | 0.006 |
| Marginal R^2^ / Conditional R^2^: 0.015 / 0.395 | | | | |

### **AICc tables and model results of effects of *F_GRM_* on telomere length**

**Table S5:** LMMs of variation in early-life telomere length predicted by *F_GRM_* (*n*=371). All models included random intercepts for brood identity and year.

| Model | | ∆AICc | df | w |
| --- | --- | --- | --- | --- |
| 1 | log(TL) = tarsus + sex + island + age + hatch day + *F_GRM_* + *F_GRM_* * sex + *F_GRM_* * island | 0 | 12 | 0.65 |
| 2 | log(TL) = tarsus + sex + island + age + hatch day + *F_GRM_* + *F_GRM_* * island | 2.1 | 11 | 0.22 |
| 3 | log(TL) = tarsus + sex + island + age + hatch day + *F_GRM_* + *F_GRM_* * sex | 3.4 | 11 | 0.12 |
| 4 | log(TL) = tarsus + sex + island + age + hatch day + *F_GRM_* | 7.5 | 10 | 0.06 |
| 5 | log(TL) = tarsus + sex + island + age + hatch day | 23 | 9 | 0.00 |

**Table S6:** LMMs of variation in early-life telomere length predicted by *F_GRM_* (*n*=371) with or without a break point at the mean *F_GRM_*=0.016. All models included random intercepts for brood identity and year.

| Model | | ∆AICc | df | w |
| --- | --- | --- | --- | --- |
| 1 | log(TL) = tarsus + sex + island + age + hatch day + *F_GRM_*<0.016 *+ F_GRM_*>0.016 + *F_GRM_*<0.016 * island + *F_GRM_*>0.016 * island | 0 | 13 | 0.62 |
| 2 | log(TL) = tarsus + sex + island + age + hatch day + *F_GRM_*<0.016 *+ F_GRM_*>0.016 + *F_GRM_*<0.016 * island + *F_GRM_*>0.016 * island + *F_GRM_*<0.016 * sex + *F_GRM_*>0.016 * sex | 3.1 | 15 | 0.13 |
| 3 | log(TL) = tarsus + sex + island + age + hatch day + *F_GRM_* + *F_GRM_* * sex + *F_GRM_* * island | 3.1 | 12 | 0.13 |
| 4 | log(TL) = tarsus + sex + island + age + hatch day + *F_GRM_* + *F_GRM_* * island | 5.3 | 11 | 0.05 |
| 5 | log(TL) = tarsus + sex + island + age + hatch day + *F_GRM_*<0.016 *+ F_GRM_*>0.016 | 6.1 | 11 | 0.03 |
| 6 | log(TL) = tarsus + sex + island + age + hatch day + *F_GRM_* + *F_GRM_* * sex | 6.6 | 11 | 0.02 |
| 7 | log(TL) = tarsus + sex + island + age + hatch day + *F_GRM_*<0.016 *+ F_GRM_*>0.016 + *F_GRM_*<0.016 * sex + *F_GRM_*>0.016 * sex | 8.9 | 13 | 0.01 |
| 8 | log(TL) = tarsus + sex + island + age + hatch day + *F_GRM_* | 10.6 | 10 | 0.00 |
| 9 | log(TL) = tarsus + sex + island + age + hatch day | 26.1 | 9 | <0.001 |

### **AICc tables and model results of effects of** ***F_ROH_* on telomere length**

**Table S7:** LMMs of variation in early-life telomere length predicted by *F_ROH_* (*n*=371). All models included random intercepts for brood identity and year.

| Model | | ∆AICc | df | w |
| --- | --- | --- | --- | --- |
| 1 | log(TL) = tarsus + sex + island + age + hatch day + *F_ROH_* + *F_ROH_* * sex | 0 | 11 | 0.31 |
| 2 | log(TL) = tarsus + sex + island + age + hatch day + *F_ROH_* | 0.2 | 10 | 0.29 |
| 3 | log(TL) = tarsus + sex + island + age + hatch day | 1.3 | 9 | 0.16 |
| 4 | log(TL) = tarsus + sex + island + age + hatch day + *F_ROH_* + *F_ROH_* * sex + *F_ROH_* * island | 1.8 | 12 | 0.12 |
| 5 | log(TL) = tarsus + sex + island + age + hatch day + *F_ROH_* + *F_ROH_* * island | 2.1 | 11 | 0.11 |

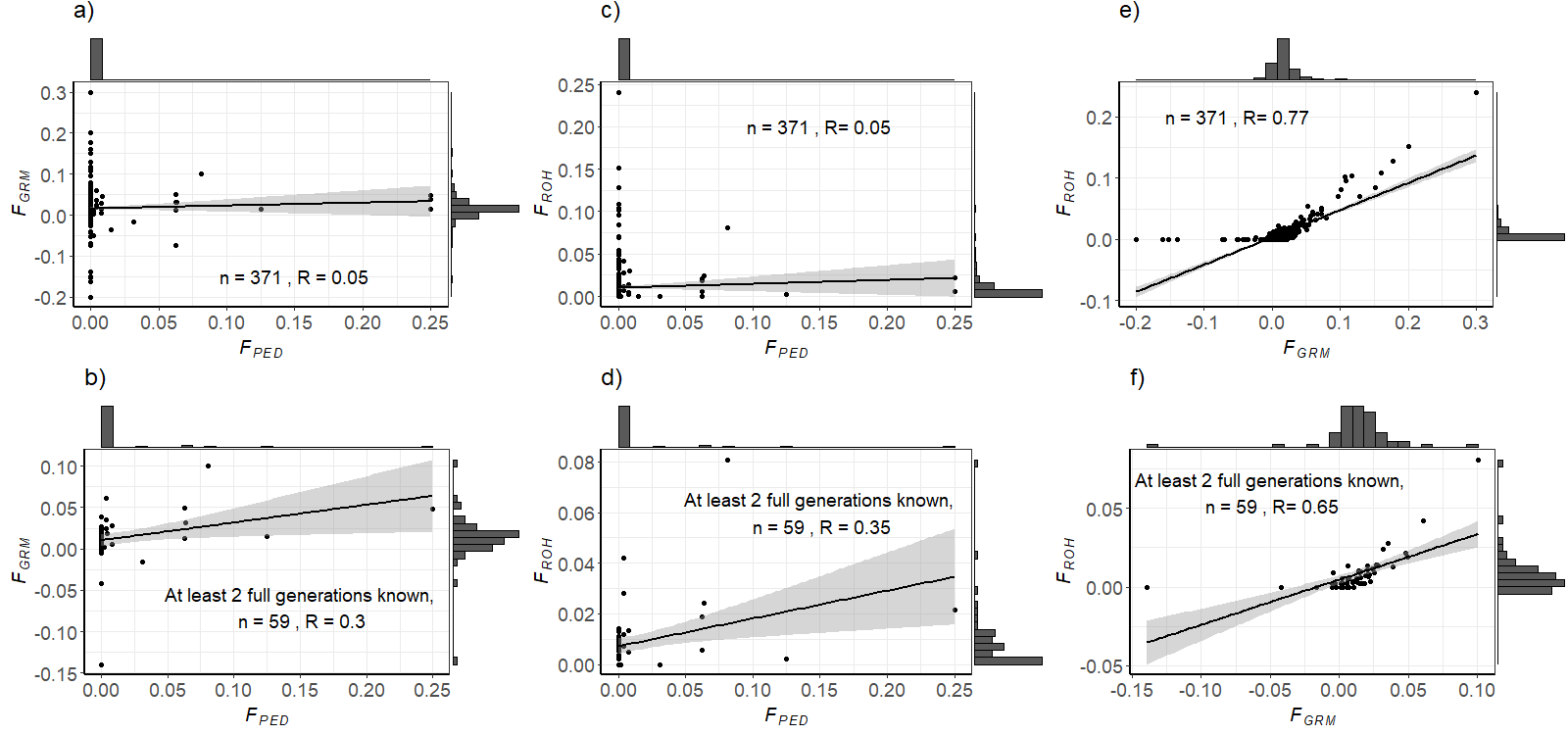

### **Correlations between different measures of inbreeding**

**Figure S1:** Sample sizes (n) and Pearson’s correlation coefficients (R) between different estimators of inbreeding. Lower panels show the same correlations restricted to individuals with at least two full ancestral generations known. Black lines are linear regression lines with 95% confidence intervals shown in grey.
